# Elongation Index as a Sensitive Measure of Cell Deformation in High-Throughput Microfluidic Systems

**DOI:** 10.1101/2020.02.17.953323

**Authors:** S. J. Hymel, H. Lan, D. B. Khismatullin

## Abstract

One of the promising approaches for high-throughput screening of cell mechanotype is microfluidic deformability cytometry (mDC) in which the apparent deformation index (DI) of the cells stretched by extensional flow at the stagnation point of a cross-slot microchannel is measured. The DI is subject to substantial measurement errors due to cell offset from the flow centerline and velocity fluctuations in inlet channels, leading to artificial widening of DI vs. cell size plots. Here, we simulated an mDC experiment using a custom computational algorithm for viscoelastic cell migration. Cell motion and deformation in a cross-slot channel was modeled for fixed or randomized values of cellular mechanical properties (diameter, shear elasticity, cortical tension) and initial cell placement, with or without sinusoidal fluctuations between the inlet velocities. Our numerical simulation indicates that mDC loses sensitivity to changes in shear elasticity when the offset distance exceeds 5 μm, and just 1% velocity fluctuation causes an 11.7% drop in the DI. The obtained relationships between the cell diameter, shear elasticity, and offset distance were used to establish a new measure of cell deformation, referred to as “Elongation Index” (EI). In the randomized study, the EI scatter plots were visibly separated for the low and high elasticity populations of cells, with a mean of 300 and 3,500 Pa, while the standard DI output was unable to distinguish between these two groups of cells. The successful suppression of the offset artefacts with a narrower data distribution was shown for the EI output of MCF-7 cells.

**Statement of Significance:** This study establishes a new measure of high-throughput microfluidic deformability cytometry, referred to as “elongation index”, that is not subject to cell offset artefacts and can sensibly and reliably detect disease-induced changes in mechanical properties of living cells.

## I. Introduction

Malignancy and hereditary blood disorders such as sickle cell disease cause reorganization of the intracellular structure that alters the ability of the cell to deform under applied stress(1–5). Various techniques have been developed to measure deformability of living cells, which can be classified into 1) single-cell methods such as micropipette aspiration and atomic force microscopy(6–13) and 2) microfluidics-based deformability cytometry(14–19). Microfluidic methods provide several advantages for disease diagnosis over traditional singlecell techniques: 1) high throughput and easy operation, 2) physiological flow conditions, 3) reduced risk of cell activation, and 4) ability to detect the stage of the cell cycle during deformability measurement. There is a large variation in mechanical properties of living cells measured by single-cell methods, even for healthy cells of the same phenotype, which can be attributed to cell activation and measurement at different stages of the cell cycle.

In microfluidic deformability cytometry (mDC), the cells are stretched by extensional flow at the stagnation point of a cross-slot microchannel at a rate up to 2,000 cells/s. This method specifically produces cell deformation index (DI) vs cell size scatter plots. The cells with different mechanical properties (“mechanotype”) are distinguished by comparing the “eyes” of these plots (most populated values of DI and size). mDC has been successfully used to identify non-activated and activated leukocytes, inflammation and malignancy in blood and pleural fluids, and stem cell pluripotency(14–17). It operates under inertial flow conditions (Reynolds number Re >10) provided the extensional stress do not rupture the cells. The threshold flow rate for cell rupture in mDC was measured by Armistead(20) and Bae(21) for different cell types in Newtonian and viscoelastic fluids.

mDC remains a pure empirical technique that does not go beyond cell size and DI measurement. DI is expected to be a function of cell size, cortical tension, and bulk viscoelasticity of the cell, but it also depends on the lateral position of the cell just before it enters the extensional flow region. The drift of the cells to a lateral position between the centerline and the channel wall always occur in mDC experiments, leading to measurement errors and, in particularly, to artificially wide distribution of the DI and cell size. Asymmetric cell stretching due to pressure fluctuations, which is caused by flow splitting, also contributes to these errors. None of the modeling studies addressed these issues. Simple analytical expressions that relate the DI with channel geometry and cell’s viscoelastic properties based on Maxwell and Kevin-Voigt models exist only for the cell moving along the flow centerline under low-velocity, Stokes flow conditions(21–24). Commercial software packages (e.g., COMSOL Multiphysics^®^) that can only simulate flow but not cell deformation in a cross-slot channel have been employed to optimize the channel geometry for mDC(20, 21, 25–27). This approach worked well for Re ≤ 10, but failed for higher Re. Data analysis based on predictive mDC models that account for DI changes due to cell offset from the centerline and flow disturbances is necessary to further improve mDC accuracy and sensitivity.

In the present study, we have used our custom computational algorithm for migration and deformation of a viscoelastic cell to simulate an mDC experiment. The DI of the cell was evaluated for different cell size, shear elasticity, cortical tension, offsets from the flow centerline in the inlet channels, and pressure fluctuations between two inlet channels. Based on this analysis, the approximate relationships between DI and cell offsets were proposed. Using these relationships in the numerical simulation where the diameter, shear elasticity, and offsets of the cell were randomized, we demonstrated that mDC sensitivity to cell mechanotype can be substantially improved. Specifically, applying this correction theory to the DI numerical data led to a much narrower distribution of DI and size for MCF-7 cells than the experimental density plots.

## II. Methods

### A. Numerical Algorithm

In this work, we have used a custom fully three-dimensional numerical algorithm for living cell migration and deformation, referred to as viscoelastic cell adhesion model (VECAM)(28, 29). In VECAM, the cell and its external environment are a multiphase continuum with moving interfaces (e.g., cell’s cortical layer) tracked by the volume-of-fluid (VOF) method. The velocity field inside and outside the cell is determined from the solution of the continuity and Navier-Stokes equations in a Marker-and-Cell (MAC) grid with the values of physical parameters for each phase averaged over a grid element. The cortical tension force per unit volume **f** is calculated in each element by the continuous surface force (CSF) method:

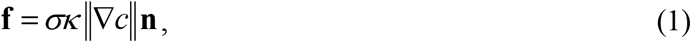

where *σ* is the cortical tension, **n** = ▿*c*/||▿*c*|| the outward unit normal to the cell surface; *κ* = –▿·**n** the local mean curvature of the cell surface; *c* = *c*(*t*, **x**) the concentration function that takes the value of 1 inside the cell, between 0 and 1 at the interface, and 0 outside the cell; and **x** = (*x,y,z*) the position vector in the Cartesian coordinate system. Since the cell’s mass density is nearly the same as the extracellular fluid density, gravity was neglected in the simulation.

The viscoelasticity of the cell cytoplasm is described by the Oldroyd-B model:

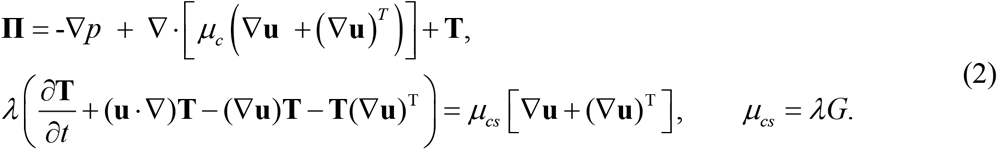

Here **Π** is the total stress tensor that includes the contributions from pressure *p*, cytosolic shear viscosity *μ_c_*, and cytoskeletal viscoelasticity determined by the extra stress tensor **T**. Other variables in (2) are the velocity vector **u** = (*u,v,w*), mass density *ρ*, cytoskeletal shear elastic modulus *G*, cytoskeletal shear viscosity *μ_cs_*, and cytoplasmic relaxation time *λ*. The cytoplasmic viscosity *μ_cp_* was defined as the sum of cytosolic and cytoskeletal shear viscosities: *μ_cp_* = *μ_c_* + *μ_cs_*. It should be noted that single-phase viscoelastic cell models have been employed for both theoretical and experimental analysis of blood cells and circulating tumor cells(1, 9, 23, 30–34).

### B. Modeling of mDC experiment

The simulation of cell deformation was performed in a computational domain with a cross-slot channel geometry (Fig. 1a). The height and width of the channel (30 μm and 60 μm, respectively)(17) and the volumetric flow rate (425 μL/min in the inflow channel)(14) matched the configuration of previous mDC experiments. The total length of the computational domain including the 120-μm-long inflow channel was 300 μm. Fully developed flow was first established in the channel. The cell was then placed at 90 μm from the cross-slot center in the flow direction (X-direction) and at the flow centerline or varying distance away from the centerline (offset distance) in the cross-section of the channel (Y- and Z-directions). The cell diameter varied from 8 μm to 26 μm. Its relaxation time was fixed at 0.17 s(1, 4) and shear elasticity ranged from 50 Pa to 20 kPa. The shear viscosity of the extracellular fluid, which modeled as a Newtonian fluid, was 0.001 Pa·s (1 cP). The cytosolic shear viscosity was equal to the extracellular fluid viscosity. The trajectories of the simulated cell motion in the cross-slot channel as well as cell shape changes at different time instants were recorded (Fig. 1b). The maximum deformation index (DI) was calculated from the cell shape data.

**Fig. 1:**
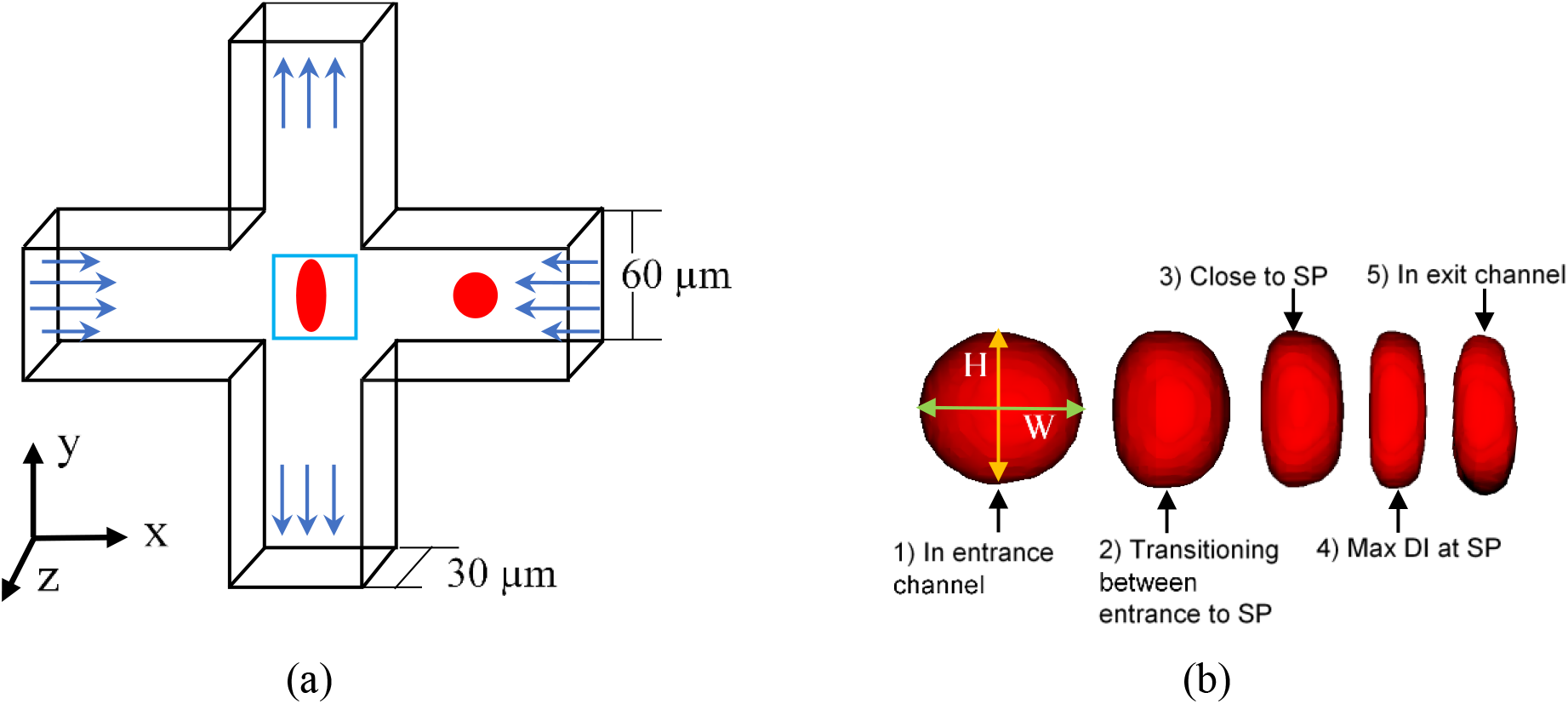
(a) Schematic of the cross-flow channel geometry with velocity profiles at the inlets and outlets. A cell initially located in one of the inflow channels flows in the cross-slot region where it experiences maximum deformation at the stagnation point (SP) as shown in the blue box. (b) Shape changes of the cell as it flows through the cross-flow channel, according to the numerical simulation. As the cell progresses through the channel, the shape is recorded by measuring the Deformation Index (DI) = major axis (H, orange) / minor axis (W, green) lengths.

Due to pressure fluctuations at two inlets of the cross-slot channel and a lateral drift of the cell away from the centerline, most of the cells are displaced from the center (stagnation point or SP) of the channel during mDC experiments. If the cell is perfectly located at the SP, it experiences equal but opposite hydrodynamic forces, which trap the cell and cause extensive cell deformation. This scenario rarely occurs in mDC. To accommodate both these conditions, the initial Y-offset distance of the cell, Y_off_, ranged in the numerical simulation from 0 to a maximum of 23 μm. The latter was selected such that to have at least 0.5 μm clearance of the cell from the wall. The initial Z-offset distance, Z_off_, was between 0 and 7 μm. To test the effect of flow fluctuations due to uneven pressures at the inlets, we imposed a sinusoidal oscillation on the centerline velocity *U_c_* at both inlets (inlets 1 and 2):

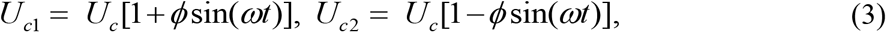

where *ϕ* is the dimensionless amplitude of velocity oscillation ranged from 0 to 0.1 (0 to 10%) in the simulation and *ω* is the angular frequency, which was equal to 21 s^-1^. The maximum variation in the centerline velocity between the inlets was 1.6 m/s.

### C. Correction of the DI data

The numerical data for cell deformation at different *Y_off_* and *Z_off_* were compared to the centerline DI data, and the resulting offset-based errors were fit into exponential relationships between the DI, cell diameter D, and the total offset distance

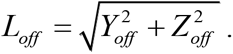

The obtained regression data for the errors were then subtracted from the original DI’s, leading to the corrected measure of cell deformation, referred to as “elongation index” (EI):

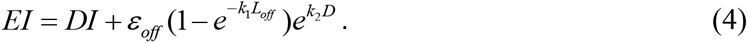

Note that the cell offset causes an artificial decrease in the DI, i.e., the offset-based errors are proportional to a negative of *ε_off_*. Regression analysis was done by GraphPad Prism (GraphPad Software, La Jolla, CA), with the coefficient of determination *R*^2^ of at least 87%.

### D. Mesh Refinement

All the data were produced on a computational mesh with cubic grid elements of 1.0 μm in size (regular mesh). To test if this mesh size provides sufficient accuracy, the simulation of single cell motion initially located at the channel centerline was additionally done on the coarse mesh (2.0 μm in size) and the fine mesh (0.5 μm). In this mesh refinement study, the cell diameter *D* was 18 μm and the shear elasticity *G* was either 1.0 or 5.0 kPa. In the case of a more deformable cell (*G* = 1.0 kPa), there was a 25% difference in DI between the coarse and fine mesh, but less than a 3% difference between the regular and fine mesh. For a less deformable cell (*G* = 5.0 kPa), only 7% difference in DI was between the coarse and fine mesh and less than a 2% difference was between the regular and fine mesh. Thus, the regular mesh simulation had less than 3% error, when compared with the fine mesh data, but required 4× less computational resources and took half the time of the fine mesh simulation.

## III. Results and Discussion

Fig. 2(a) shows the fully developed flow field in a cross-slot channel obtained numerically in the absence of the cell. Red color indicates peak velocity achieved and blue color designates the zero-velocity areas, which includes the SP region in the middle of the computational domain. To test if our computational model can reproduce the experimental data on HL-60 cell deformation and breakup in a cross-slot channel(20), we first performed the numerical simulation of a 12 μm-diameter cell with shear elasticity *G* = 100 Pa. The cell was initially located at the inlet channel centerline and exposed to inlet flow rate from 67.5 to 675 μL/min, corresponding to Re from 25 and 250. Note that the total flow rate is twice the inlet flow rate. The numerical DI data (red circles in Fig. 2b) agreed closely with the experimental data by Armistead *et al*.(20) (blue squares) in which only the cells moving along the flow centerline were taken into account. In further analysis, Re was fixed at 157, which was less than the threshold value (180) for cell breakup(20) and agreed with experimental conditions in Gossett *et al*.(14)

**Fig. 2:**
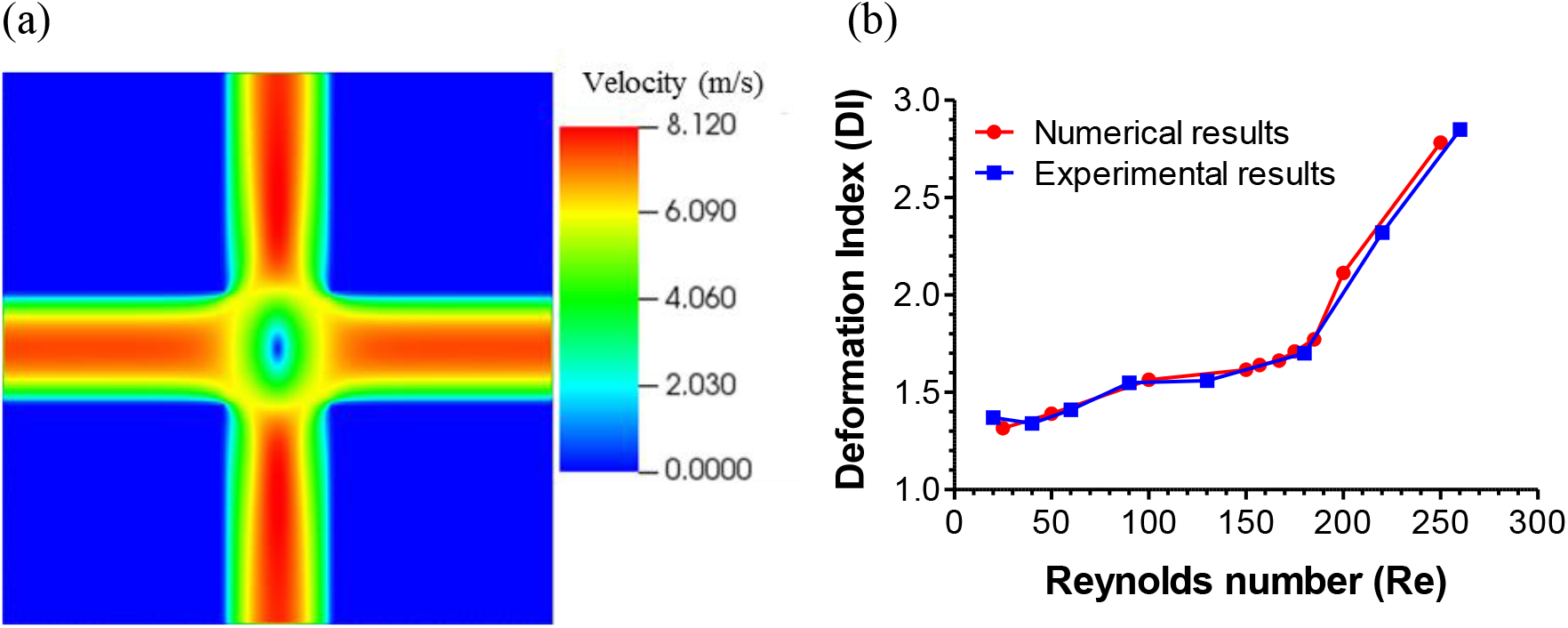
(a) Fully developed flow profile through the mid-plane of the cross-slot channel without the cell present, according to the numerical simulation. (b) Comparison of the experimental (blue squares) and numerical relationships (red circles) between DI and Re for HL-60 cells. In (b), the numerical cell size and shear elasticity were 12 μm and 100 Pa, respectively.

Fig. 3 illustrates how the DI changes with cell size and elastic properties (shear elasticity, cortical tension) provided the cell was originally located at the inlet channel centerline. A bigger cell experienced larger deformation in the SP area even in the absence of changes in shear elasticity (Fig. 3a). The DI rapidly decreased when shear elasticity changed from 50 Pa to 1 kPa and slowly plateaued at shear elasticity above 10 kPa (Fig. 3a). Cortical tension had a significant effect on the DI at low shear elasticity (up to 1 kPa) and a nominal effect at higher elasticity (Fig. 3b). At *G* = 100 Pa, the DI peaked at 4.9 at σ = 300 pN/μm but decreased to a minimum of 3.6 at 3,000 pN/μm. When *G* = 1 kPa, there was a steady decrease in the DI from 3.6 to 3.2 with σ between 30 and 3,000 pN/μm.

**Fig. 3:**
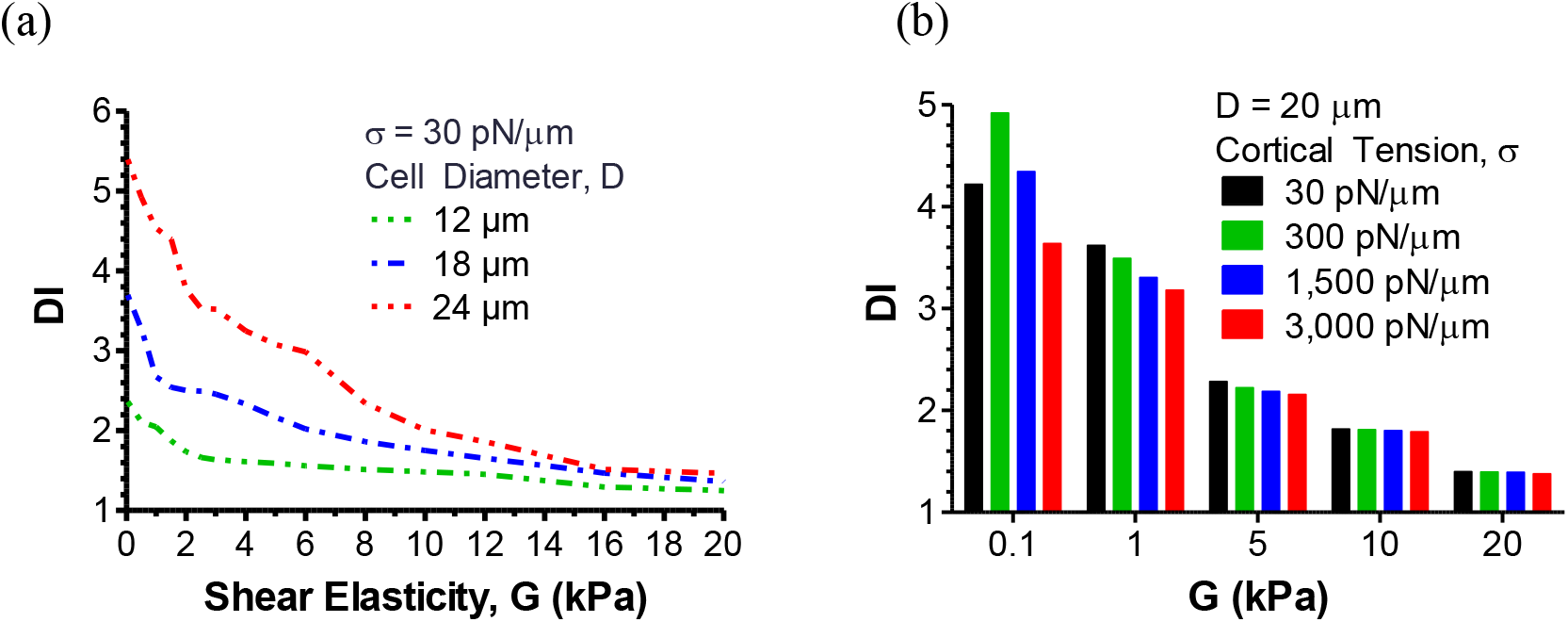
(a) DI vs cytoskeletal shear elasticity for 12, 18, and 24 μm-diameter cells, according to the numerical simulation. (b) DI vs cytoskeletal shear elasticity of a 20 μm cell for different values of the cortical tension.

The data shown in Fig. 4 point out that the wide spread of DI and cell sizes in mDC experiments (Fig. 4a) is due to the lateral and transverse displacements of the cell from the centerline (Y- and Z-offsets, respectively). The DI rapidly dropped when the Y-offset (Y_off_) increased from 0.1 to 1.0 μm, and this effect became more pronounced with an increase in cell size (Fig. 4b, c). For instance, the DI of a 24-μm cell with shear elasticity of 1 kPa decreased from 4.23 at Y_off_ of 0.1 μm to 2.97 at Y_off_ of 1.0 μm (30% difference, Fig. 4b). The DI reached the plateau at Y_off_ of ~5 μm at which the sensitivity of this parameter to shear elasticity was lost (Fig. 4c). Even without lateral offset, mDC cannot distinguish the cells with shear elasticities between 500 Pa and 10 kPa when the Z-offset (Z_off_) becomes 2.0 μm or higher (Fig. 4d). At Z_off_ greater than 2.0 μm, there was less than a 5% difference in the DI between the cells with shear elasticity of 50 Pa and 10 kPa. This indicates that mDC at large Z_off_ is not sensitive to elasticity change within three orders of magnitude. Much more drastic changes in the DI were seen when the cell was offset in both lateral and transverse directions (Fig. 4e). An 18-μm cell with shear elasticity of 1 kPa had the DI of 3.22 at Y_off_ of 0.1 μm and no Z_off_ (Fig. 4b) or 2.47 at Z_off_ of 0.1 μm and no Y_off_ (Fig. 4e), which was then decreased to 1.42 at Y_off_ of 1.0 μm and Z_off_ of 2.0 μm (44% difference; Fig. 4e). These data also show that the effect of Z_off_ on the DI reduces with an increase in Y_off_. When the Y_off_ is above 5.0 μm, the transverse displacement of the cells contributes insignificantly to errors of mDC measurement, in comparison to their lateral displacement.

**Fig. 4:**
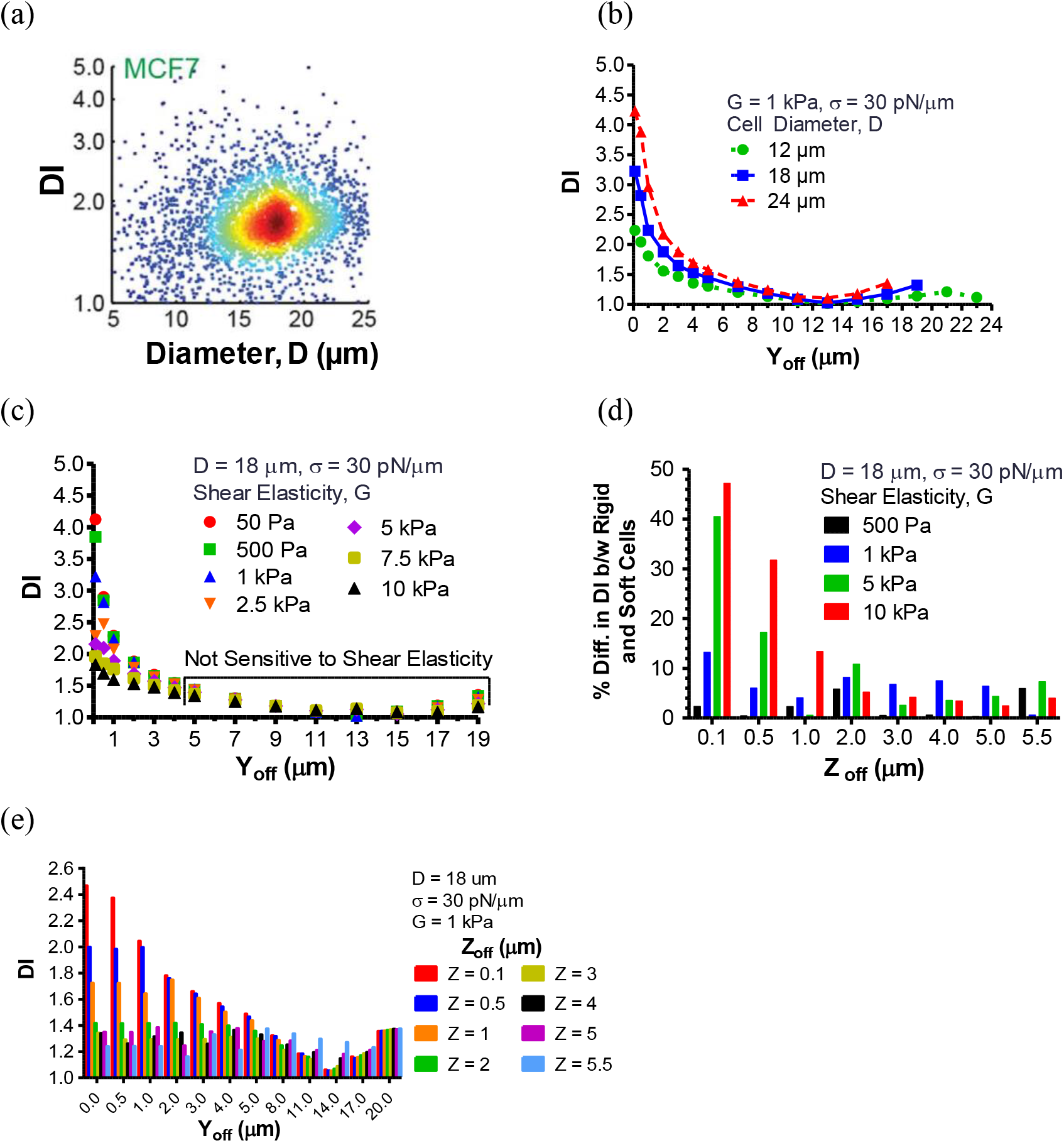
(a) Density plot of deformability index vs diameter of MCF-7 cells generated by mDC (Reproduced from Fig. 5d in Masaeli *et al.* with permission). (b) DI vs. Y_off_ for the cell with shear elasticity of 1 kPa and different cell diameters. (c) DI vs. Y_off_ for the 18 μm-diameter cells with different values of the cytoskeletal shear elasticity. (d) The effect of Z_off_ on the percent difference in DI between the rigid cells (*G* ≥ 500 Pa) and the soft cell *(G* = 50 Pa). Y_off_ was 0 μm. (e) DI vs. Y_off_ for the 18 μm-diameter cell initially located at different Z_off_.

To assess errors in DI measurement resulting from inlet flow perturbations, we studied numerically cell motion and deformation in a cross-slot channel when the centerline velocity between the inlets oscillated with amplitude *φ* = 0, 1, 3, 5 and 10%. As seen in Fig. 5(a), there was a significant deviation in the cell trajectory from the symmetric flow case (*ϕ* = 0%, black solid line) even when the oscillation amplitude was 1% (blue dashed line). With this small variation, the cell traveled longer to the cross-slot SP than the cell in symmetric flow. A further increase in *φ* led to the bounce-back effect (rise and drop in the total distance; cf. green, red and purple dotted lines) at which the cell moved past the SP and then returned to it through decaying oscillation in lateral (Y-) direction. This behavior has been already observed in mDC experiments. Fig. 5(b) illustrates that the shape of the cell trapped in the SP drastically changed with *φ.* When compared to symmetric flow, the DI decreased by 11.7, 19.2, 29.6, and 35.2% for *φ* = 1, 3, 5 and 10%, respectively (Fig. 5c). Inlet flow perturbations also caused the cell to drift from the centerline in the transverse (Z-) direction.

**Fig. 5:**
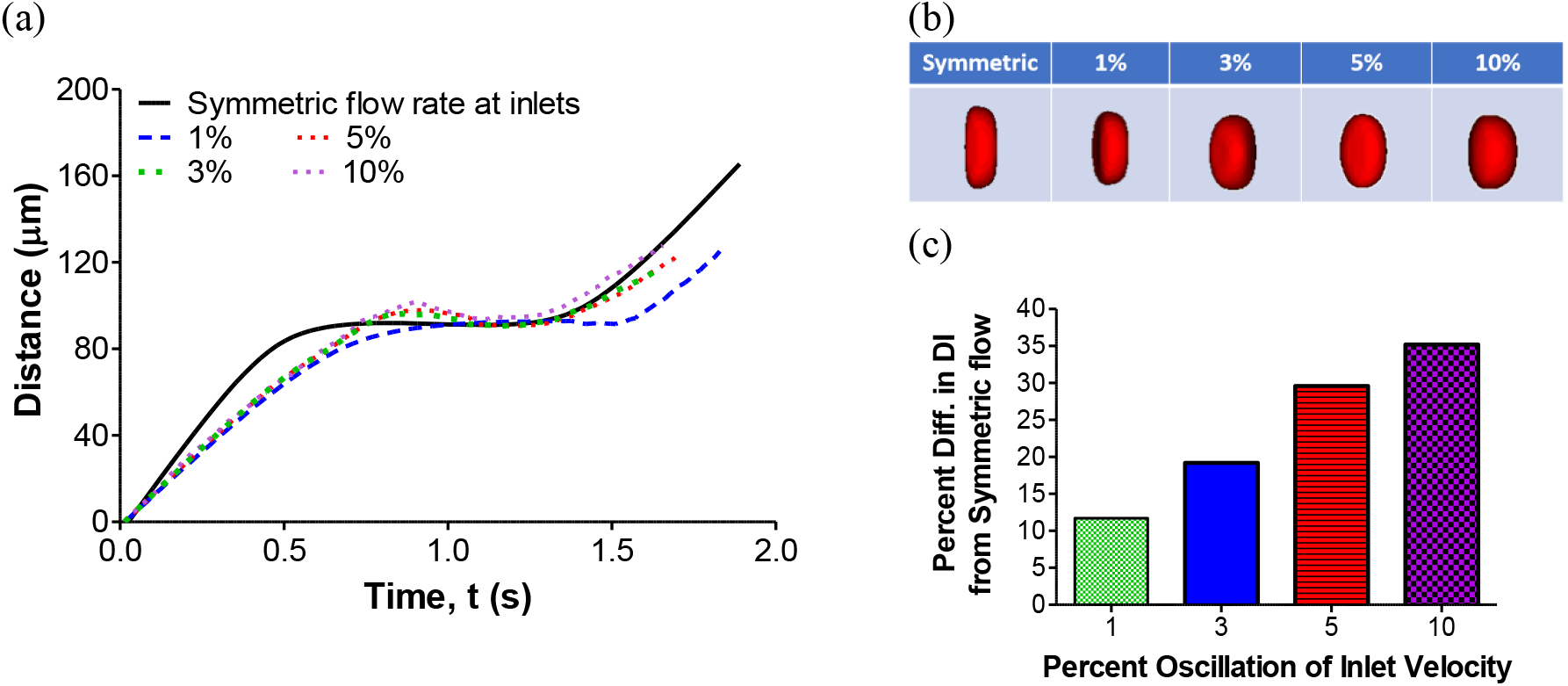
The numerical effect of oscillatory perturbation in the inlet velocity on cell motion and deformation in a cross-slot channel. (a) Total distance traveled by the cell and (b) cell shape at the SP region of the cross-slot channel for different amplitudes of sinusoidal flow oscillation between the inlets. (c) Percent difference in the cell DI between symmetric flow (no fluctuation between the inlets) and disturbed flow due to velocity fluctuations. The diameter and elasticity of the cell were 18 μm and 1 kPa, respectively.

To account for variability in mechanical properties of the cells and their location in the inlet channel during mDC experiment, we conducted the numerical study of cell deformation in a cross-slot channel at which the initial cell placement, cell size, and shear elasticity were randomized via pseudo-random normal sampling. In this randomization study, the cell diameter D changed from 10 to 24 μm, Y_off_ ranged 0.1 and 10 μm, and Z_off_ was between 0 and 2.5 μm. Fig. 6(a) shows the DI vs cell diameter scatter plots for two populations of cells: 1) low elasticity group (mean *G* = 300 Pa, blue dots) and 2) high elasticity group (mean *G* = 3.5 kPa, red dots). The scatter plots for the same sampling of cell size and elasticity but with a very small offset in the lateral direction (Y_off_ = 0.1 μm) and no offset in the transverse direction (Z_off_ = 0 μm) are displayed in Fig. 6(b). The latter study was done to get the DI values for the cell moving along the inlet channel centerline. (Note that we had to place a small Y-offset to avoid complete trapping of the cell within the SP region.) From comparison between Figs. 6(a) and 6(b), it is clearly seen that the lateral and transverse offsets led to artificial enlargement of the DI output to the point when it was no longer possible to distinguish between the cells in the low and high elasticity groups that have more than a 10× difference in shear elasticity.

**Fig. 6:**
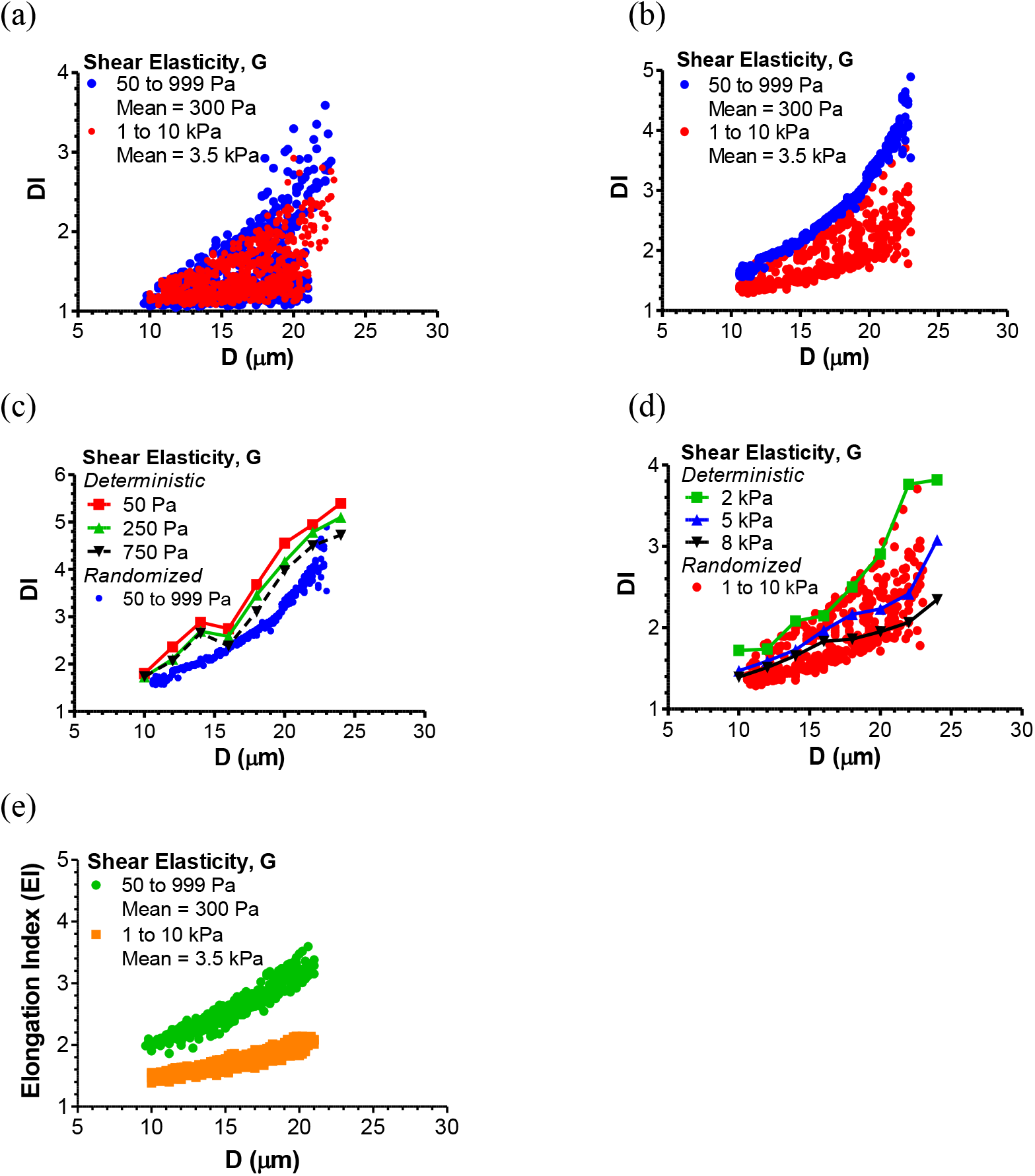
Scatter plots for cell deformation vs. cell diameter for low and high elasticity groups before and after data correction by (4). (a-d) DI vs. D plots, according to the numerical model with normal random distribution of *D* and *G* and uniform random distribution of Y_off_ and Z_off_ (a) or fixed minimal offset (Y_off_ = 0.1 μm and Z_off_ = 0 μm, b-d). In (c) and (d), minimum-offset randomized data for the low (c) and high elasticity groups (d) were compared with the deterministic results obtained for the cell with different shear elasticity initially located at the inlet channel centerline. (e) EI vs. D distribution obtained by correcting the offset errors in the original DI data (a) via (4). There was a clear separation of the EI between the low and high elasticity groups, which was not seen in the DI data. The shear elasticity of the cells in the low elasticity group (blue) ranged from 50 to 999 Pa, with a mean of 300 Pa and a standard deviation of 200 Pa. The high elasticity group (red) had the shear elasticity between 1 and 10 kPa, with a mean of 3.5 kPa and a standard deviation of 2 kPa. The cell diameter changed from 10 to 24 μm, with a mean of 17 μm and a standard deviation of 4 μm. The ranges for Y_off_ and Z_off_ in the randomized study were 0.1 to 10 μm and 0.1 to 2.5 μm, respectively.

When the cell offset was minimal (Fig. 6b), randomized data for the low elasticity group slightly under-predicted the deterministic DI values for the cell moving exactly at the centerline (Fig. 6c). The high elasticity group had a wider distribution of DI’s at minimal offset (Fig. 6b), but all the DI data fell within the deterministic centerline values (Fig. 6d).

With applying (4) to the randomized data in Fig. 6a, the errors due to offsets had been minimized, and the resulting EI scatter plots were clearly separated for the low and high elasticity groups (Fig. 6e). This indicates that the sensitivity of mDC to shear elasticity can be substantially improved by correcting the DI output by (4).

The effect of shear elasticity on regression parameters *ε_off_, k*_1_, and *k*_2_ in (4) are shown in Fig. 7. For the EI output in Fig. 6(e), *ε_off_* was 0.395 and 0.274, *k* was 1.007 and 0.412 μm^-1^, and *k*_2_ was 0.071 and 0.057 μm^-1^ for the low and high elasticity groups, respectively. It should be noted that the EI does not have the same dependence on cell diameter, as predicted by the centerline data. It rather represents a central tendency for the minimum offset DI distribution in Fig. 6(b).

**Fig. 7:**
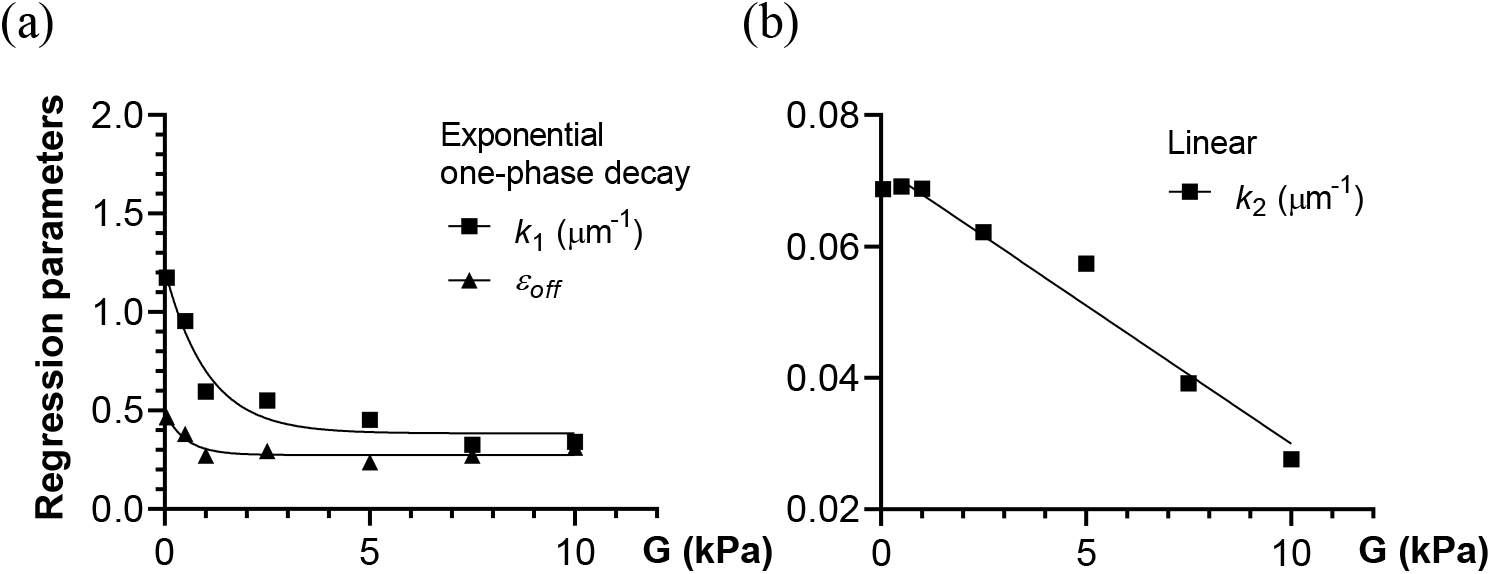
Dependence of parameters in regression (4) on shear elasticity. (a) *ε_off_* and *k*_1_ vs. *G* obtained from the deterministic study (symbols, cf. Fig. 4e) and fitting those data to the exponential one-phase decay model (lines). (b) *k*_2_ vs. *G* for the deterministic study (squares, cf. Fig. 4b) and its fitting to the linear model (line).

In the last numerical study with randomized values of parameters, we generated the DI values for MCF-7 noninvasive breast cancer cells. The mean shear elasticity and diameter of these cells were selected to be 413 Pa and 18 μm, based on previous measurements(35, 36). Fig. 8 shows the distribution of the DI vs. cell diameter with uniform random offsets (blue dots) and minimum fixed offset (black) as well as the EI vs cell diameter output (green) obtained by correcting the DI values (blue) by (4). The following values of regression parameters was used in this study: *ε_off_* = 0.372, *k*_1_ = 0.945 μm^-1^, and *k*_2_ = 0.070 μm^-1^. The correction of DI values by (4) led to a narrower, compact distribution of the cell deformation data. The resulting EI values had the same trend with cell diameter as the minimum-offset DI, thus indicating successful suppression of the offset errors.

**Fig. 8:**
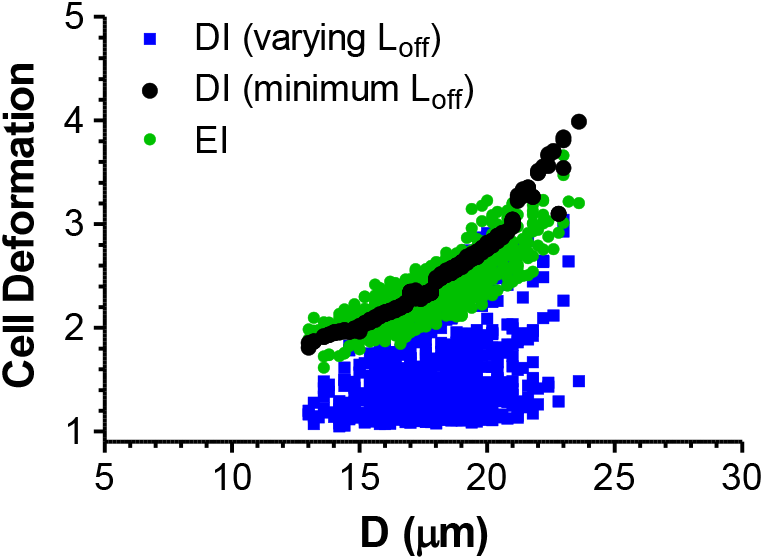
Numerical scatter plots for MCF-7 cell deformation vs. diameter before (blue, black) and after offset error correction (green). The cell diameter *D* and shear elasticity *G* were normally distributed and the offsets (Y_off_ and Z_off_) were either uniformly distributed (blue) or fixed at a minimal value (Y_off_ = 0.1 μm and Z_off_ = 0 μm, black). The shear elasticity for MCF-7 cells ranged from 200 to 600 Pa, with a mean of 413 Pa and a standard deviation of 100 Pa. The cell diameter range was from 13 to 24 μm, with a mean of 18 μm and a standard deviation of 2 μm. Y_off_ and Z_off_ in the fully randomized analysis (blue) varied from 0.1 to 10 μm and 0.1 to 2.5 μm, respectively.

## IV. Conclusion

Cross-slot microfluidic deformability cytometry (mDC) is a promising high-throughput approach for mechanotypic screening of living cells, but it is plagued with measurement errors associated with cell offset from the flow centerline(24, 37) and pressure or velocity fluctuations at the inlet channels. One of the ways to reduce offset errors is to eliminate the off-distance deformation index (DI) data(20, 23), which can make mDC less powerful in assessing changes in cell mechanotype. We have demonstrated via predictive computational modeling that the mDC output can be sensitized to mechanical properties of the cells such as shear elasticity without removing viable off-distance data. In this new approach, the DI is replaced with a new measure, referred to as “Elongation Index” (EI), which accounts for offset errors. The EI formula (Eq. 4) has been derived based on regression analysis of the numerical data for cell deformation in a cross-slot microchannel obtained for different values of cell shear elasticity, diameter, and initial offset from the inlet channel centerline. This formula requires the knowledge of the offset distance and cell diameter, both of which can be measured from acquired images of cells. We also have quantified the effect of flow-splitting-induced fluctuations in inlet velocities on DI measurement. In particular, 1% velocity fluctuation was found to cause an 11.7% drop in the DI. This error further increased to 35.2% at 10% fluctuation. Overall, the present study demonstrates that mDC can be significantly improved by its integration with predictive computational models of deformable cell migration such as VECAM.

## Author Contributions

S.J.H. and D.B.K designed this study and wrote the manuscript. H.L. and S.J.H. developed the computational algorithm. S.J.H. performed numerical analysis of cell deformation in a cross-slot microfluidic channel.

## Acknowledgments

The authors acknowledge funding from National Science Foundation (grants No. 1301286 and 1438537) and Louisiana Board of Regents (LEQSF(2011-14)-RD-A-24). This research was supported in part using high performance computing (HPC) resources and services provided by Technology Services at Tulane University, New Orleans, Louisiana. This work also used the Bridges system, which is supported by National Science Foundation grant number ACI-1445606, at the Pittsburgh Supercomputing Center (PSC). The authors thank Hideki Fujioka and Zhongyi Sheng for help with computational work and Dino Di Carlo for helpful discussion.

## References

1. Khismatullin, D. B. 2009. The Cytoskeleton and Deformability of White Blood Cells. Curr. Top. Membr. 64:47–111.

2. Seltmann, K., A. W. Fritsch, J. A. Kas, and T. M. Magin. 2013. Keratins significantly contribute to cell stiffness and impact invasive behavior. P. Natl. Acad. Sci. USA 110(46):18507–18512.

3. Coughlin, M. F., D. R. Bielenberg, G. Lenormand, M. Marinkovic, C. G. Waghorne, B. R. Zetter, and J. J. Fredberg. 2013. Cytoskeletal stiffness, friction, and fluidity of cancer cell lines with different metastatic potential. Clin. Exp. Metastasis 30(3):237–250.

4. Tomaiuolo, G. 2014. Biomechanical properties of red blood cells in health and disease towards microfluidics. Biomicrofluidics 8(5).

5. Ellett, F., J. Jorgensen, A. L. Marand, Y. M. Liu, M. M. Martinez, V. Sein, K. L. Butler, J. Lee, and D. Irimia. 2018. Diagnosis of sepsis from a drop of blood by measurement of spontaneous neutrophil motility in a microfluidic assay. Nat. Biomed. Eng. 2(4):207–214.

6. Shojaei-Baghini, E., Y. Zheng, and Y. Sun. 2013. Automated micropipette aspiration of single cells. Ann. Biomed. Eng. 41(6):1208–1216.

7. Bento, D., R. O. Rodrigues, V. Faustino, D. Pinho, C. S. Fernandes, A. I. Pereira, V. Garcia, J. M. Miranda, and R. Lima. 2018. Deformation of Red Blood Cells, Air Bubbles, and Droplets in Microfluidic Devices: Flow Visualizations and Measurements. Micromachines-Basel 9(4).

8. Hochmuth, R. M. 2000. Micropipette aspiration of living cells. J. Biomech. 33(1):15–22.

9. Dong, C., R. Skalak, K. L. Sung, G. W. Schmid-Schonbein, and S. Chien. 1988. Passive deformation analysis of human leukocytes. J. Biomech. Eng. 110(1):27–36.

10. Li, Q. S., G. Y. H. Lee, C. N. Ong, and C. T. Lim. 2008. AFM indentation study of breast cancer cells. Biochem. Bioph. Res. Co. 374(4):609–613.

11. Costa, K. D. 2003. Single-cell elastography: Probing for disease with the atomic force microscope. Dis. Markers 19(2-3):139–154.

12. Lee, L. M., and A. P. Liu. 2014. The Application of Micropipette Aspiration in Molecular Mechanics of Single Cells. J. Nanotechnol. Eng. Med. 5(4):0408011–0408016.

13. Dulinska, I., M. Targosz, W. Strojny, M. Lekka, P. Czuba, W. Balwierz, and M. Szymonski. 2006. Stiffness of normal and pathological erythrocytes studied by means of atomic force microscopy. J. Biochem. Bioph. Meth. 66(1-3):1–11.

14. Gossett, D. R., H. T. K. Tse, S. A. Lee, Y. Ying, A. G. Lindgren, O. O. Yang, J. Y. Rao, A. T. Clark, and D. Di Carlo. 2012. Hydrodynamic stretching of single cells for large population mechanical phenotyping. P. Natl. Acad. Sci. USA 109(20):7630–7635.

15. Tse, H. T. K., D. R. Gossett, Y. S. Moon, M. Masaeli, M. Sohsman, Y. Ying, K. Mislick, R. P. Adams, J. Y. Rao, and D. Di Carlo. 2013. Quantitative Diagnosis of Malignant Pleural Effusions by Single-Cell Mechanophenotyping. Sci. Transl. Med. 5(212).

16. Masaeli, M., D. Gupta, S. O’Byrne, H. T. K. Tse, D. R. Gossett, P. Tseng, A. S. Utada, H. J. Jung, S. Young, A. T. Clark, and D. Di Carlo. 2016. Multiparameter mechanical and morphometric screening of cells. Sci. Rep. 6.

17. Lin, J., D. Kim, H. T. Tse, P. Tseng, L. L. Peng, M. Dhar, S. Karumbayaram, and D. Di Carlo. 2017. High-throughput physical phenotyping of cell differentiation. Microsyst. Nanoeng. 3.

18. Nyberg, K. D., K. H. Hu, S. H. Kleinman, D. B. Khismatullin, M. J. Butte, and A. C. Rowat. 2017. Quantitative Deformability Cytometry: Rapid, Calibrated Measurements of Cell Mechanical Properties. Biophys. J. 113(7):1574–1584.

19. Gill, N. K., C. Ly, K. D. Nyberg, L. Lee, D. P. Qi, B. Tofig, M. Reis-Sobreiro, O. Dorigo, J. Y. Rao, R. Wiedemeyer, B. Karlan, K. Lawrenson, M. R. Freeman, R. Damoiseaux, and A. C. Rowat. 2019. A scalable filtration method for high throughput screening based on cell deformability. Lab Chip 19(2):343–357.

20. Armistead, F. J., J. G. De Pablo, H. Gadelha, S. A. Peyman, and S. D. Evans. 2019. Cells Under Stress: An Inertial-Shear Microfluidic Determination of Cell Behavior. Biophys. J. 116(6):1127–1135.

21. Bae, Y. B., H. K. Jang, T. H. Shin, G. Phukan, T. T. Tran, G. Lee, W. R. Hwang, and J. M. Kim. 2016. Microfluidic assessment of mechanical cell damage by extensional stress. Lab Chip 16(1):96–103.

22. Tanyeri, M., E. M. Johnson-Chavarria, and C. M. Schroeder. 2010. Hydrodynamic trap for single particles and cells. Appl. Phys. Lett. 96(22).

23. Guillou, L., J. B. Dahl, J. G. Lin, A. I. Barakat, J. Husson, S. J. Muller, and S. Kumar. 2016. Measuring Cell Viscoelastic Properties Using a Microfluidic Extensional Flow Device. Biophys. J. 111(9):2039–2050.

24. Henon, Y., G. J. Sheard, and A. Fouras. 2014. Erythrocyte deformation in a microfluidic cross-slot channel. Rsc Adv. 4(68):36079–36088.

25. Galindo-Rosales, F. J., M. S. N. Oliveira, and M. A. Alves. 2014. Optimized cross-slot microdevices for homogeneous extension. Rsc Adv. 4(15):7799–7804.

26. Haward, S. J., M. S. N. Oliveira, M. A. Alves, and G. H. McKinley. 2012. Optimized Cross-Slot Flow Geometry for Microfluidic Extensional Rheometry. Phys. Rev. Lett. 109(12).

27. Kim, J., J. Lee, C. Wu, S. Nam, D. Di Carlo, and W. Lee. 2016. Inertial focusing in non-rectangular cross-section microchannels and manipulation of accessible focusing positions. Lab Chip 16(6):992–1001.

28. Khismatullin, D. B., and G. A. Truskey. 2005. Three-dimensional numerical simulation of receptor-mediated leukocyte adhesion to surfaces: Effects of cell deformability and viscoelasticity. Phys. Fluids 17(3).

29. Khismatullin, D. B., and G. A. Truskey. 2012. Leukocyte rolling on P-selectin: a three-dimensional numerical study of the effect of cytoplasmic viscosity. Biophys. J. 102(8):1757–1766.

30. Lim, C. T., E. H. Zhou, and S. T. Quek. 2006. Mechanical models for living cells - A review. J. Biomech. 39(2):195–216.

31. Zhou, C., P. Yue, and J. J. Feng. 2007. Simulation of neutrophil deformation and transport in capillaries using newtonian and viscoelastic drop models. Ann. Biomed. Eng. 35(5):766–780.

32. Takeishi, N., Y. Imai, T. Yamaguchi, and T. Ishikawa. 2015. Flow of a circulating tumor cell and red blood cells in microvessels. Phys. Rev. E 92(6).

33. Bronkhorst, P. J. H., G. J. Streekstra, J. Grimbergen, E. J. Nijhof, J. J. Sixma, and G. J. Brakenhoff. 1995. A new method to study shape recovery of red blood cells using multiple optical trapping. Biophys. J. 69(5):1666–1673.

34. Evans, E., and A. Yeung. 1989. Apparent Viscosity and Cortical Tension of Blood Granulocytes Determined by Micropipet Aspiration. Biophys. J. 56(1):151–160.

35. Lekka, M., D. Gil, K. Pogoda, J. Dulinska-Litewka, R. Jach, J. Gostek, O. Klymenko, S. Prauzner-Bechcicki, Z. Stachura, J. Wiltowska-Zuber, K. Okon, and P. Laidler. 2012. Cancer cell detection in tissue sections using AFM. Arch. Biochem. Biophys. 518(2):151–156.

36. Arya, S. K., K. C. Lee, D. Bin Dah’alan, Daniel, and A. R. A. Rahman. 2012. Breast tumor cell detection at single cell resolution using an electrochemical impedance technique. Lab Chip 12(13):2362–2368.

37. Cha, S., T. Shin, S. S. Lee, W. Shim, G. Lee, S. J. Lee, Y. Kim, and J. M. Kim. 2012. Cell Stretching Measurement Utilizing Viscoelastic Particle Focusing. Anal. Chem. 84(23):10471–10477.

